# Construction of a liquid-liquid phase separation system from the gel-sol transition of elongated protein microgels in a crowding agent

**DOI:** 10.1101/2020.12.08.416867

**Authors:** Yufan Xu, Runzhang Qi, Hongjia Zhu, Bing Li, Yi Shen, Georg Krainer, David Klenerman, Tuomas P. J. Knowles

## Abstract

Liquid proteinaceous materials have been frequently found in cells or tissues and are crucial for various biological processes. Unlike their solid-state counterparts, liquid-state protein compartments are challenging to engineer and control at the microscale. Conventionally, gelation (sol-gel transition) of biological molecules has been thought to be the intermediate step between liquid-liquid phase separation (LLPS) states and insoluble aggregates that are related to protein functions, malfunctions and even diseases. However, the opposite process, i.e., the gel-sol transition of materials, has not been broadly explored. Here we describe a thermoresponsive gel-sol transition of a protein in a crowded environment that results in a demixed LLPS state, contradicting the common consequence of a one-phase protein solution by the end of such transition at elevated temperature without crowding agents. We also demonstrate a simple method to monitor the gel-sol transition by showing that elongated gelatin microgels can evolve towards a spherical morphology in the crowding agents because of interfacial tension. The LLPS system was explored for the diffusion of small particles for drug-release application scenarios. Our results demonstrate a route for the rapid construction of LLPS models, where the gel-sol transition of the protein-rich phase is monitorable. The models are featured with tunable size and dimensional monodispersity of dispersed condensates. The present study can be employed in biophysics and bioengineering with practices such as 3D printing and temperature sensing.

## Introduction

Phase transitions are common guiding principles for the processing of soft materials for biological and medical studies (1–4). To date, sol-gel transitions have been substantially studied and utilised for the fabrication of solid-state materials like soft hydrogels for diverse purposes such as tissue engineering and pathology studies (1–8). Recently, there is an increasing interest in liquid proteinaceous materials, as they are found to be highly related to diverse biological functions or malfunctions for healthcare and diseases (9–12). The development of liquid-state materials is therefore a crucial advancement to functional and intelligent materials which can be generated from gel-sol transition that however is rarely studied. Liquid-state and gel-state proteins can involve different physicochemical mechanisms, and liquid-state materials are a key category of substances mimicking the native states of proteins in vivo as an important supplement of conventional solid or gel materials. Protein-rich phases in liquid-liquid phase separation (LLPS) systems have been increasingly of scientific intertest, as the proteins in relatively independent spatial domains at the microscale play key roles in the understanding of the formation of membranless organelles, silk fibres, as well as insoluble aggregates accountable for neurodegeneration (10; 11; 13–15).

LLPS systems are the precursors of the protein gels or aggregates, but the opposite process (dissolution or gel-sol transition) has not been well established (16; 17). One obstacle of the gel-sol transition in LLPS is to screen a protein material that has reversible sol-gel transition properties, and the other obstacle is to generate LLPS systems with controllable size and dispersity of the disperse phases. Factors such as pH, salt, temperature, ionic strength, and crowding agents can impact LLPS (17). Protein-rich phases from existing LLPS systems arose from the concentration of materials in solutions; the spontaneous formation of protein-rich phases has similar mechanisms of nucleation and growth, and thus the monodispersity of the LLPS is difficult to achieve. LLPS can contribute to intracellular activities that are involved with protein condensates and membraneless organelles, but extracellular proteins have not been well studied for LLPS yet (18–20). Phase separation of extracellular matrix (ECM) proteins can be taken for generic mechanisms for the assembly of proteins into fibrillar or polymeric assemblies (21). Understanding the LLPS of ECM protein, such as elastin and collagen, can contribute to the studies in aberrant assembly of proteins which can imply pathological consequences and fundamental principles for the optimisation of materials and drugs (21). The adoption of extracellular proteins or their substitutes as one phase of all-aqueous emulsions holds great potential for the rapid construction of LLPS systems for fundamental physicochemical research and medical applications in drug release and disease models.

Another challenge that LLPS studies face is the lack of fast and simple characterisation to confirm the liquidity of the condensates. Existing techniques like fluorescence recovery after photobleaching, optical tweezering, and live cell imaging usually require complex equipment and facilities (11; 18; 22; 23). The fusion of condensates has been employed for the proof of the liquidity (24), but the spontaneous contacting of condensates is stochastic. An easily manipulable method to highlight the liquid nature of materials at the microscale is therefore demanded for LLPS characterisations.

Here we describe a two-step engineering approach to the construction of a LLPS system by facilitating the thermoresponsive gel-sol transition of gelatin in a crowding agent, resulting in a demixed all-aqueous state. Firstly, gelatin microgels through physical crosslinking were engineered on a microfluidic device followed by demulsification into water. Secondly, the gelatin microgels underwent a gel-sol transition in crowding agents to produce a demixed LLPS system at elevated temperature. In contrast, a homogeneous solution was acquired with the absence of crowding agents. Monodisperse protein-rich phases can be achieved by adjusting the population of microgels. We show that the heat-triggered shape change of the elongated microgels and the fusion of the protein-rich phases can both highlight the liquidity of the protein-rich phases. As a control, enzymatically crosslinked microgels were thermostable and did not transition to liquid phase. The LLPS were further discovered dependent on the concentrations of the crowding agents and could act as a platform for the diffusion of nanoparticles for nanomedicine. The LLPS systems developed in this present study can shed light on the phase-separation and phase-transition behaviours and more practically on 3D printing and drug release for regenerative medicine.

## Results and Discussion

We used a microfluidic setup in combination with heating accessories at 37 °C to keep the gelatin solution in its liquid state as previously reported (**Figure 1**) (4; 25). A microfluidic device with a flow-focusing junction was used to generate the microrods (elongated microgels); there are two inlets and one outlet in the microfluidic device. The inlet tubing for the disperse phase (gelatin solution) was heated up to 37 °C; the inlet tubing for the continuous phase (oil) was placed at room temperature (RT; 25 °C); the outlet tubing for microdroplets in oil was placed under an ice bag. The gelatin microdroplets were generated at the microfluidic junction, and they were elongated and physically crosslinked in the thin outlet tubing. These monodisperse microrods in oil were then demulsified into water at RT or lower temperature (**Figure 1, 2c, 2d**). There was slight swelling of the microrods during the demulsification because of the water uptake. The demulsification of the monodisperse microrods allows the formation of monodisperse all-aqueous systems. Further biological and pharmaceutical studies can thus be performed with the oil-free hydrogels or hydrosols.

**Figure 1.**
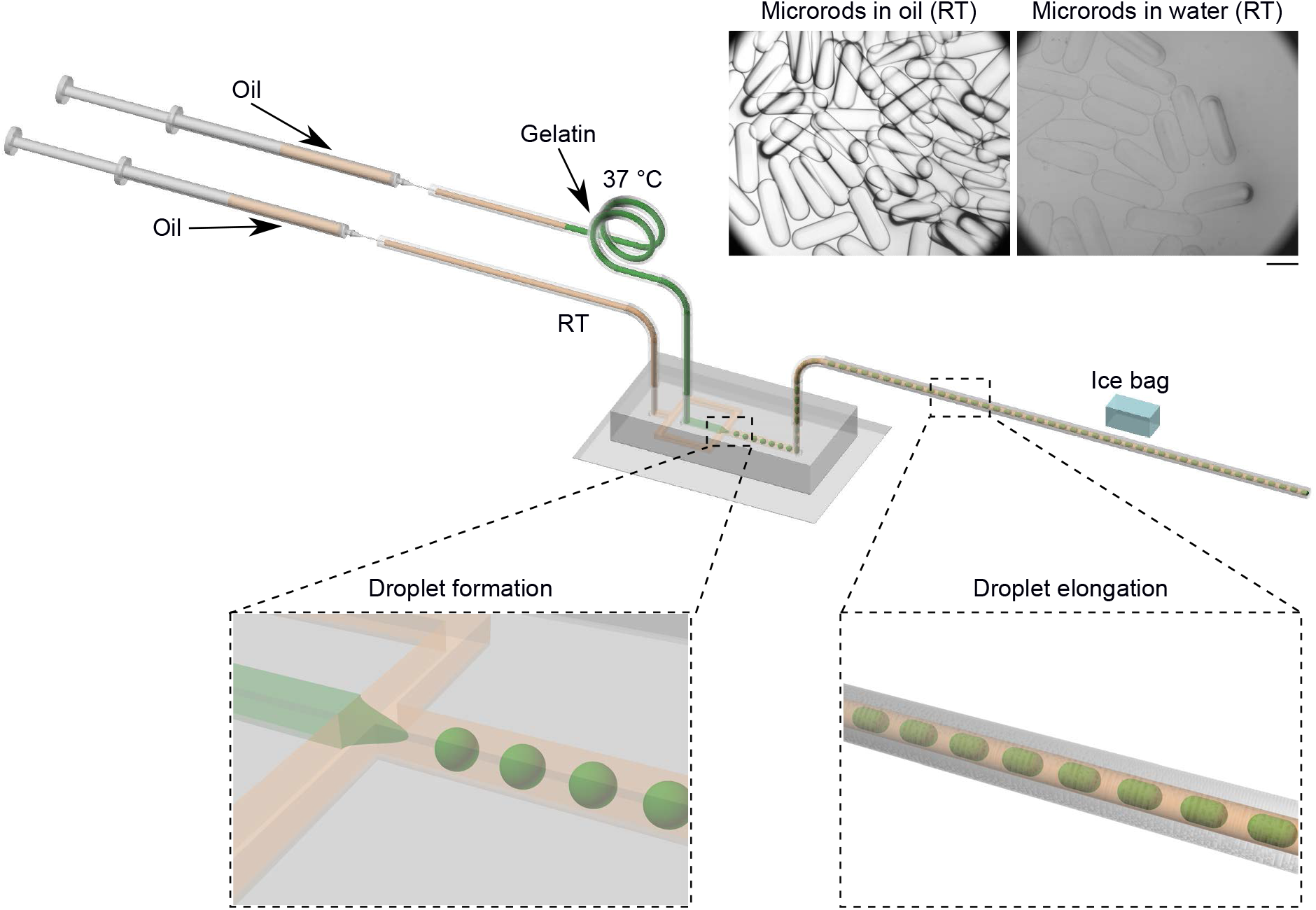
Generation of physically crosslinked protein microrods with a microfluidic approach. An inlet tubing for the gelatin solution was heated up to 37 ° C; an inlet tubing for the oil phase was placed at room temperature (RT; 25 °C); an outlet tubing was placed under an ice bag for the physical crosslinking of the elongated gelatin microdroplets in oil. The microscopy images at top right show the elongated microdroplets (microrods) in oil and in water. Scale bar, 500 μm.

**Figure 2.**
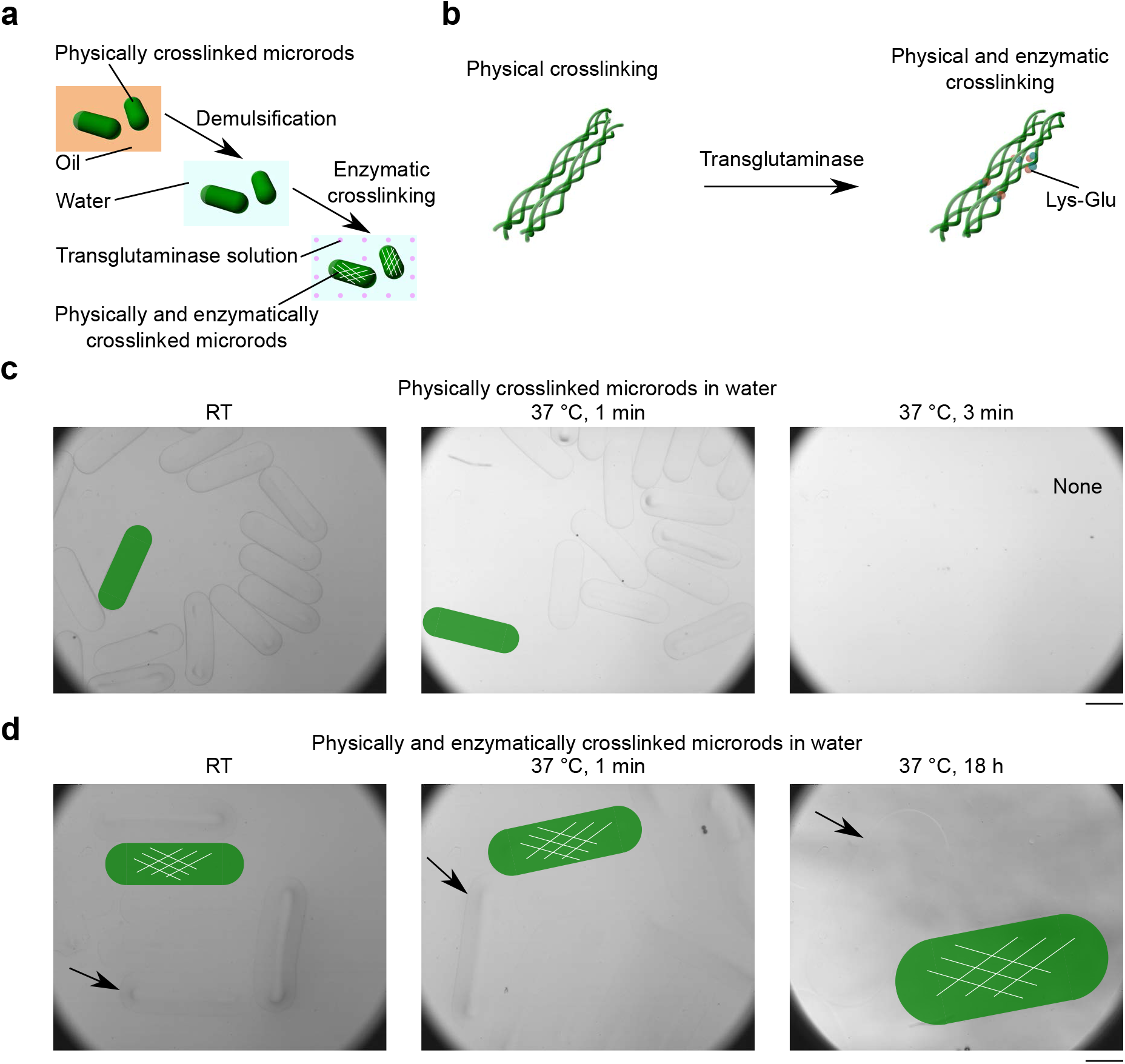
Thermostability of the microrods. a, Schematic of the physical and enzymatic crosslinking of the protein microrods. b, Molecular conformation of gelatin under different crosslinking regimes. Lys, lysine. Glu, glutamine. c, Dissolution of physically crosslinked microrods in water. Scale bar, 500 μm. d, Thermostability of physically and enzymatically crosslinked microrods in water. Arrows point to gel-water interfaces. Scale bar, 500 μm.

Gelatin and its derivatives have been widely used as the analogues of collagen that is a major component of ECM proteins (4; 26–29). Triggered by temperature, gelatin can undergo reversible sol-gel transitions, and it can be processed at physiological temperature and neutral pH at multiple scales (4; 25). Gelatin is also a protein that inherits cell adhesive motifs such as Arg-Gly-Asp from native collagen (4; 30). It is of crucial advancement to fabricate all-aqueous systems with gelatin as collagen substitutes for artificial and miniaturised ECM platforms for tissue regeneration, pathology, and drug screening or testing.

The physical crosslinking of the gelatin took place quickly as a result of cooling down in the outlet tubing (**Figure 1**). At higher temperature (e.g. 37 °C), the gelatin molecules were in the conformation of random coils and the gelatin solution was flowable in the microfluidic channels and tubings (**Figure 1**) (4; 25; 26). After cooling down (e.g. RT or lower temperature), these random coils transformed into triple helices and the gelatin became physically crosslinked hydrogels (**Figure 2b, 2c**) (4; 25; 26). The microrods dissolved at 37 °C in several minutes, as a result of the weak and reversible physical crosslinking from intermolecular forces such as hydrogen bonds, van der Waals forces, electrostatic or hydrophobic interactions (**Figure 2c**) (31–33). The elongated microgels with reversible phase transition capabilities in this present study complement previous thermostable protein assemblies such as gelatin methacrylate microrods, silk microcylinders, and elongated droplets containing amyloid fibrils (29; 34; 35).

We next developed physically and enzymatically crosslinked microrods, as a control study of the physically crosslinked microrods. Transglutaminase solution was used to enzymatically crosslink the physically crosslinked microrods from their surfaces (**Figure 2a**). Transglutaminase catalyses the formation of covalent N e-(γ-glutamyl) lysine amide bonds between the gelatin strands to form a permanent network of polypeptides, and thus the gelatin triple helices were further connected (**Figure 2b, 2d**) (4; 25; 26). Therefore, the physically any enzymatically crosslinked microrods were thermostable when heated up to 37 °C (**Figure 2d**), which is different from the physically crosslinked microgels (Figure 2c). The thermostable microrods are promising candidates for implants *in vivo* as well as drug carriers at physiological temperature *in vitro* (4; 25).

We further extended the application of the physically crosslinked microrods to construct LLPS models, by mixing gelatin microrods with PEG solution at RT followed by heating up the mixture to 37 °C (**Figure 3ai**). Physically crosslinked microrods stayed elongated in the microrod-PEG system at RT, which means the microrods were still at the gel state at RT (**Figure 3c**). These microrods became spherical when heated up to 37 °C, which highlights the liquefaction of gelatin microrods driven by PEG-gelatin liquid-liquid interfacial tension (**Figure 3c**). The liquefaction agrees with the degradability of the physically crosslinked microrods (**Figure 2c**). As a control, the physically and enzymatically crosslinked microrods demonstrated no obvious deformation at 37 °C (**Figure 3d**). The physically and enzymatically gelatin were at gel state both at RT and 37 °C, which is in line with the thermostability of the microrods by robust covalent bonds (**Figure 2d**).

**Figure 3.**
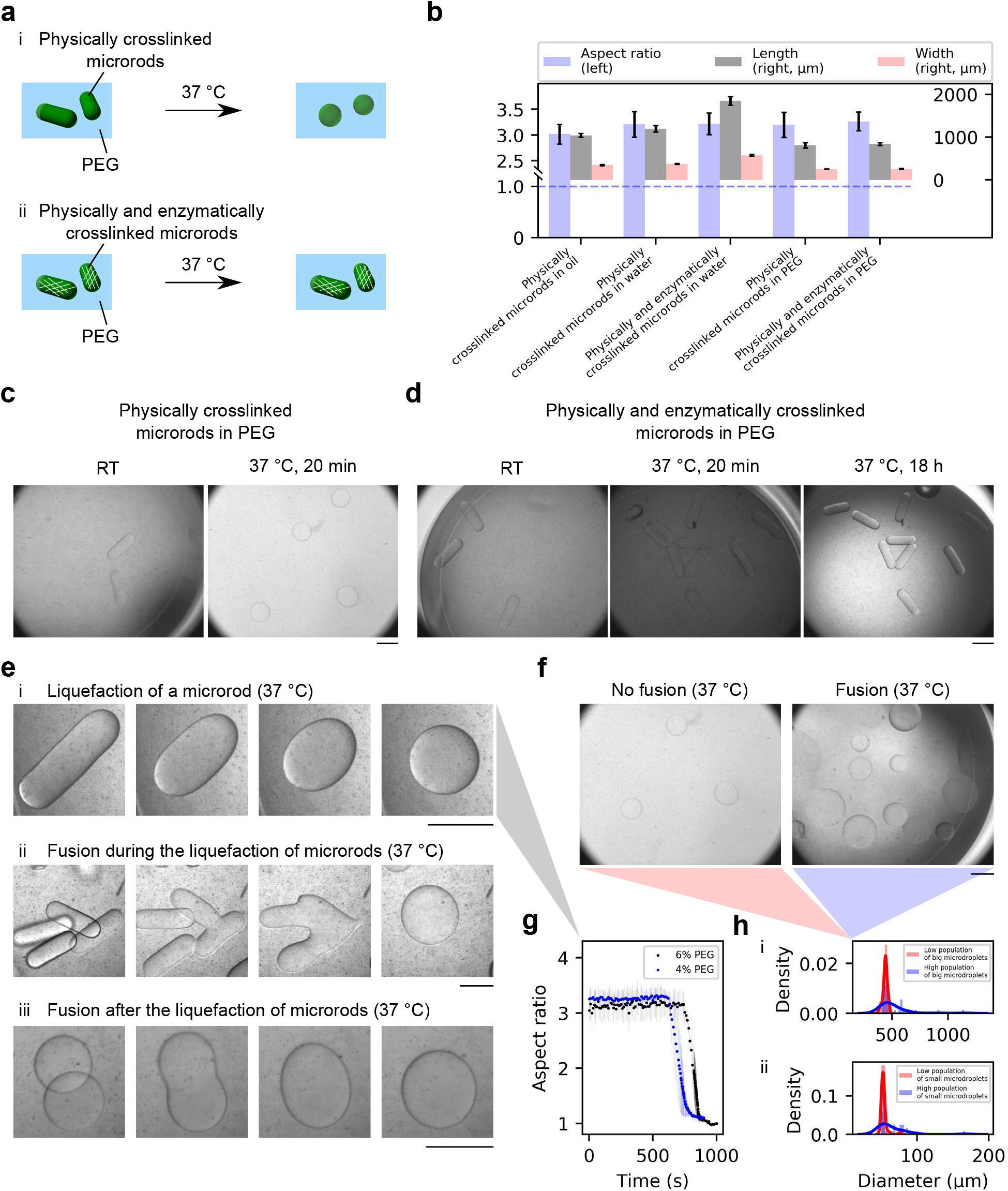
Gel-sol to sol-sol transition of gelatin-PEG all-aqueous systems. a, Schematics of the morphologies of the microrods in the gelatin-PEG all-aqueous systems before and after heating up. i, Physically crosslinked microrods. ii, Physically and enzymatically crosslinked microrods. b, Geometrical characterisation (left axis, aspect ratio; right axis, length and width) of protein microrods in oil, water, and PEG solution at RT. Sample size for each, 30. The blue dashed line indicates the aspect ratio of a spherical microdroplet. c, Physically crosslinked microrods became spherical in PEG solution at 37 °C. Scale bar, 500 μm. d, Physically and enzymatically crosslinked microrods maintained elongated in PEG solution at 37 °C. Scale bar, 500 μm. Continued on the following page. Figure 3 e, Liquefaction of physically crosslinked protein microrods (Movies available). i, Liquefaction of a single microrod. ii, Fusion during the liquefaction of several microrods. iii, Fusion after the liquefaction of two microrods. Scale bar, 500 μm. f, Fusion studies with low (left) and high (right) population of physically crosslinked protein microrods in PEG solution after incubation at 37 °C. g, Evolution of the aspect ratio of microrods during liquefaction (ei). h, Monodisperse and polydisperse LLPS from big microgels and small microgels. i, Big microdroplets are shown in f (Figure S1i). Sample sizes, 38 (low microgel population) and 39 (high microgel population). ii, Small microdroplets are shown in Figure S1ii. Sample sizes, 40 (low microgel population) and 40 (high microgel population).

The liquefaction of an individual microrod (physically crosslinked) was recorded in a thermostatic chamber (**Figure 3ei**). It took about ten minutes to trigger the deformation of spherification, and the major deformation lasted for approximately two minutes (**Figure 3ei, 3g**). The heat transfer is a key factor for the time evolution, as it took time to heat up the container, PEG solution, and the microrods. The liquefied gelatin microdroplets could fuse when they were close enough (**Figure 3eiii**). This fusion can take place when one droplet sank onto another droplet that was already at the bottom of the container, or when several droplets near each other were already at the bottom of the container. Fusion was also found during the liquefaction of several microrods (**Figure 3eii**).

All-aqueous emulsions are promising models with compartmentalised protein domains, as these oil-free emulsions can be used to perform many biological processes that take place in water (12; 36). Conventional approaches of producing all-aqueous emulsions include bulk mixing, electrospray, and 3D printing (12; 36–40). However, it is challenging to produce monodisperse LLPS emulsions by bulk mixing or 3D printing (37–39). Electrospray can produce monodisperse LLPS emulsions, but would require extra electric fields (36; 40). In order to manipulate the dispersity of the LLPS system, the fusion can be avoided or utilised. It is more likely to get a more monodisperse LLPS system with low microgel population in the PEG solution (**Figure 3f, 3hi**). Higher population density of microrods increases the possibility that microrods touched each other or were close enough to each other. Bulk mixing of gelatin solution and PEG solution would lead to polydisperse LLPS emulsion (39). This present study, in contrast, proposes a controllable approach to making monodisperse LLPS emulsion by adjusting the population density of gelatin microrods. This approach can be applied to spherical microgels as well. The dimension of the condensates can be readily adjusted by the incorporation of microgels with a wide range of sizes (**Figure 3h, S1**).

We presented the geometrical data of gelatin microrods in different continuous phases (**Figure 3b**). The aspect ratio of gelatin microrods increased during demulsification (3.02, physically crosslinked microrods in oil; 3.21, physically crosslinked microrods in water; 3.22, physically and enzymatically crosslinked microrods in water); the aspect ratio maintained similar when transferred from water into PEG solution (3.20, physically crosslinked microrods in PEG; 3.26, physically and enzymatically crosslinked microrods in PEG) (**Figure 3b**). It is possible that the microrods in oil are more compressed by the gel-oil interfacial tension in the longitudinal direction than the width direction, because oil-gel interfacial tension can be much bigger than water-gel interfacial tension (36). During the demulsification, there was a higher resilience in the longitudinal direction, which increased the aspect ratio. The microrods swelled when demulsified into water, followed by a further swelling during the enzymatic crosslinking; this implied that physically crosslinked gelatin and physically and enzymatically crosslinked gelatin have different capabilities of water uptake (**Figure 3b**). The microrods then shrank or dehydrated in PEG solution as a result of osmosis (**Figure 3b**). The dehydration was enhanced with ascending PEG concentrations (**Figure 4a**). These results show that the gelatin microrods were responsive to the external environments, and the concept is promising for soft and smart building blocks of complex structures for sensing applications (41; 42).

**Figure 4.**
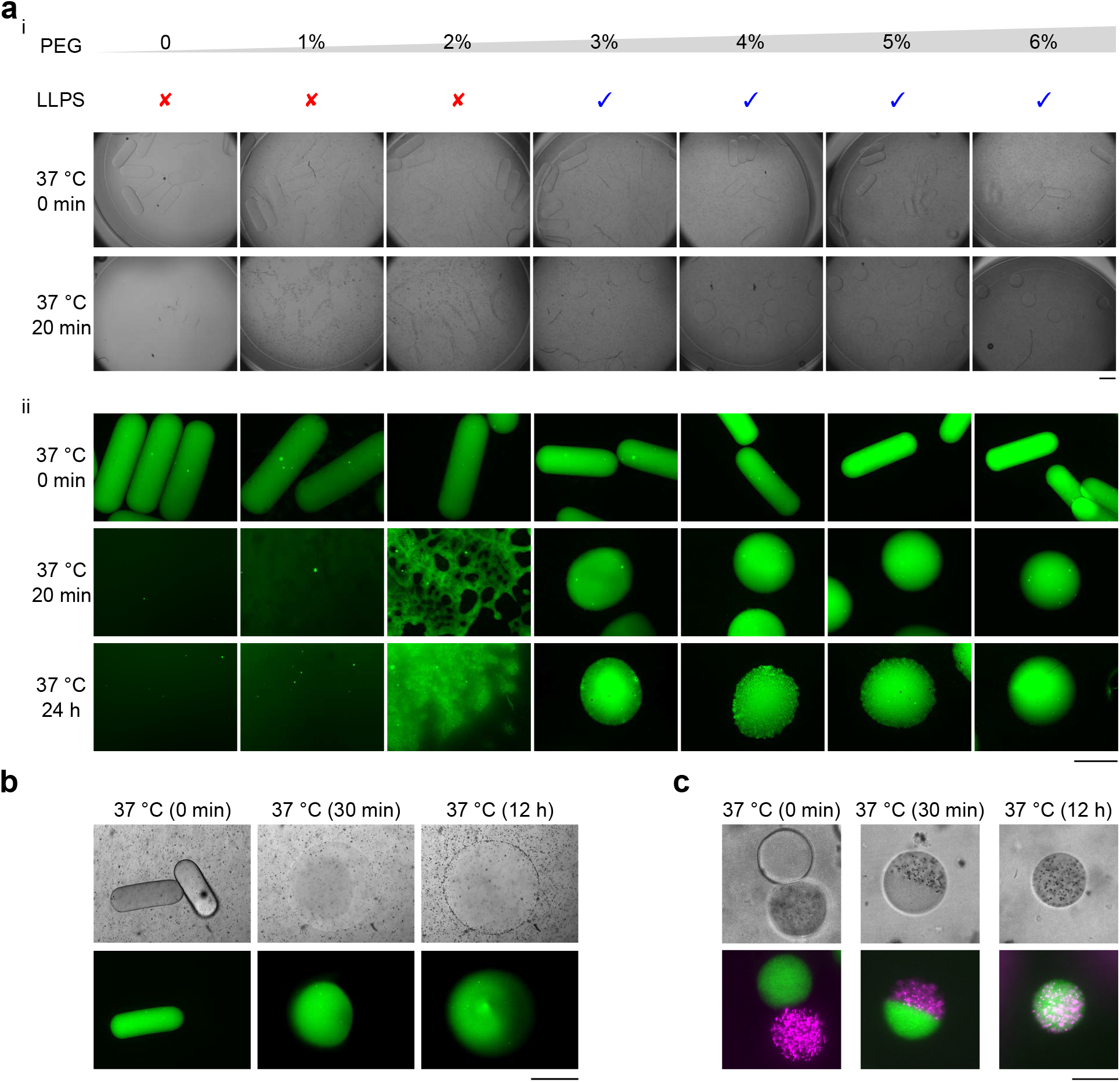
LLPS and nanosphere diffusion. a, LLPS with increasing PEG concentration. i, Brightfield microscopy. ii, Fluorescent microscopy. Scale bar, 500 μm. b, Diffusion of green nanospheres from a nanosphere-laden droplet to a nanosphere-free droplet at 37 °C (See Figure S2iii). Scale bar, 500 μm. c, Mutual diffusion of green nanospheres and red microspheres between nano/microsphere-laden droplets at 37 °C (See Figure S3iii). Scale bar, 50 μm.

We found that LLPS depended on the PEG concentrations; after heating the microrod-PEG systems, spherical gelatin-rich phases emerged in PEG of high concentrations (3%–6%) (**Figure 4ai**). Green nanospheres were premixed in the gelatin microrods, and fluorescence microscopy further highlighted this finding (**Figure 4aii**). Wetting of the gelatin-rich phase was found in the LLPS in 2% PEG (**Figure 4aii**), and no obvious LLPS was found in PEG of lower concentration (0 or 1%) (**Figure 4ai, 4aii**). PEG solutions of higher concentration have a higher depletion effect to support the LLPS and the protein condensates, which agrees with previous studies (25; 39).

We studied the diffusion of nanospheres in the LLPS system, which can inspire the drug release and diffusion in nanomedicine therapies. It was a noticeable phenomenon that the fluorescent nanospheres diffused from the gelatin-rich phase to the PEG-rich phase in PEG of low concentration (0%–2%) (**Figure 4aii**), while this diffusion was not obvious in PEG of higher concentrations (3%–6%). This can be explained by the higher diffusion coefficient of nanospheres in less viscous PEG solutions. Upon fusion, migration of nanospheres from one protein-rich droplet to the other protein-rich droplet was also observed (**Figure 4b**). The gelatin-rich microdroplets fused when heated up; the boundary of the droplets (A, with nanospheres; B, without nanospheres) was sharp at the beginning, but turned blurred in the end, which indicated the diffusion of nanospheres between droplets (**Figure 4b, S2iii**). We then explored the mutual diffusion of nanospheres and microspheres in smaller microgels (**Figure 4c, S3iii**). A relatively complete diffusion was noticed at 37 °C in 12 h, while the spheres barely migrated at RT (**Figure S2,S3**). This finding can be applied as a new pathway for the production of Janus particles or droplets (**Figure 4c,S3ii,S3iii**).

## Conclusions

In this study we have suggested a possible route for the rapid construction of an all-aqueous system using phase transition and separation of ECM protein substitutes. We developed LLPS models with tunable dispersity of elongated thermosensitive microgels. The liquefaction of these gelatin microrods was demonstrated by the morphological variations in PEG solution, further supported by their fusion behaviours. Enzymatic crosslinking as a control underpins our findings. The LLPS was dependant on the concentration of the continuous phase and can promote the elucidation of drug release and diffusion for nanomedicine. This study could have crucial implications in tissue engineering and biophysical studies for healthcare and diseases.

## Materials and Methods

### Generation of physically crosslinked microrods

A gelatin solution (w/v, 10%) and an oil continuous phase were made as previously reported (4). A hydrophobic microfluidic device (i.e. a droplet maker containing two inlets and one outlet) was made by softlithography technique (43). An inlet tubing containing the gelatin solution was heated up to 37 °C, and an inlet tubing was placed at room temperature (RT; 25 °C) (4); an outlet tubing was cooled down by an ice bag. A digital neMESYS pump system (CETONI GmbH, Korbussen, Germany) was used to inject the gelatin solution and the oil phase into the microfluidic device. The microdroplets were formed at the flow-focusing junction of the microfluidic device at 37 °C, and then elongated and physically crosslinked in the outlet tubing covered by the ice bag. The elongated microgels (microrods) were demulsified with 10% 1H,1H,2H,2H-perfluoro-1-octanol at RT or lower temperature. 1H,1H,2H,2H-perfluoro-1-octanol was kept in fumehood. Physically crosslinked microrods were stored in water.

### Generation of physically and enzymatically crosslinked microrods

A transglutaminase solution (w/v, 2%) was made by mixing transglutaminase powder (Special Ingredients Ltd, Chesterfield, UK; product of Spain) in Milli-Q water at RT (4; 25). The physically crosslinked microrods were soaked in the transglutaminase solution at RT for 6 h. Then, Milli-Q water was used to exchange the transglutaminase solution. Physically and enzymatically crosslinked microrods were stored in water.

### Degradation of the microrods

The physically crosslinked microrods and the physically and enzymatically crosslinked microrods in a 96-well plate were heated up to 37 °C. Bright-field images were taken with a high-speed camera (MotionBLITZ EoSens Mini1-1 MC1370, Mikrotron, Unterschleissheim, Germany) on a microscope (Oberver.A1, Axio, Zeiss, Oberkochen, Germany).

### Gel-sol to sol-sol transition

A polyethylene glycol (PEG) solution (w/v, 6%) was made by dissolving PEG powder (Molecular weight 300,000; Sigma-Aldrich Co Ltd, MO, US; product of USA) in Milli-Q water at 50 °C with magnetic stirring for 8 h (25). The physically crosslinked microrods and the physically and enzymatically crosslinked microrods were respectively mixed in the PEG solution at RT, and the final PEG concentration was 5.4% (w/v). The microrod-PEG mixtures were heated up to 37 °C in a 96-well plate; bright-field images were taken with the above-mentioned high-speed camera and microscope. Videos were taken on a microscope in a thermostatic chamber at 37 °C.

### LLPS depending on PEG concentrations

PEG solutions with increasing concentrations (0%, 1%, 2%, 3%, 4%, 5%, and 6%) were used to mix gelatin microrods. The final PEG concentrations in the mixtures were 0%, 0.9%, 1.8%, 2.7%, 3.6%, 4.5%, and 5.4%. The mixtures were heated up to 37 °C in 96-well plates. 1) Bright-field images were taken with a high-speed camera (MotionBLITZ EoSens Mini1-1 MC1370, Mikrotron, Unterschleissheim, Germany) on a microscope (Oberver.A1, Axio, Zeiss, Oberkochen, Germany). 2) For another experiments, green nanospheres (200 nm, 1% solids, Fluoro-Max, Thermo Scientific, CA, US) were premixed in the gelatin (1:20, v/v). Fluorescent images of the microgels were taken with a CCD camera (CoolSNAP MYO, Photometrics, AZ, US) on a microscope (Oberver.A1, Axio, Zeiss, Oberkochen, Germany); a 49001 filter (excitation wavelength 426–446 nm, emission wavelength 460–500 nm) was used with a compact light source (HXP 120 V, Leistungselektronik Jena GmbH, Jena, Germany) (4).

### Diffusion of nanospheres between droplets

1) Two categories of microrods were made by the above-mentioned methods, one with green nanospheres and the other without green nanospheres. The two categories of microrods were mixed at 1:1 ratio and then mixed with 6% PEG solution. The PEG concentration was 5.4% in the final mixture. The mixture was heated up to 37 °C in a 96-well plate. Brightfield and fluorescent images were taken with the above-mentioned imaging facilities. 2) For mutual diffusion studies, green nanospheres (200 nm) and red microspheres (1 μm) were respectively encapsulated in spherical microgels (v/v, 1:20). These spherical microgels were made smaller than the microrods at a microfluidic flow-focusing junction, and no ice bag was used. Fluorescent images of the microgels were taken with a CCD camera (CoolSNAP MYO, Photometrics, AZ, US) on a microscope (Oberver.A1, Axio, Zeiss, Oberkochen, Germany); a 49001 filter (excitation wavelength 426–446 nm, emission wavelength 460–500 nm) and a 49004 filter (excitation wavelength 532–557 nm, and emission wavelength 570–640 nm) were, respectively, used with a compact light source (HXP 120 V, Leistungselektronik Jena GmbH, Jena, Germany) (4).

## Author contribution

Y.X. and R.Q. contributed equally to this project. Y.X. and T.P.J.K. conceived and designed the project. Y.X., R.Q., and H.Z. performed the experiments. B.L. and D.K. assisted in the imaging in Figure 3ei,ii. Y.X. and R.Q. analysed the data. Y.S. and G.K. provided advice on the project. Y.X. wrote the manuscript, and all authors commented on the manuscript.

## Acknowledgements

The research leading to these results had received funding from the Cambridge Trust (Y.X.; B.L.), the Jardine Foundation (Y.X.), Trinity College Cambridge (Y.X.), Trinity College Krishnan-Ang Studentship (R.Q.), the Honorary Trinity-Henry Barlow Scholarship (R.Q.), the China Scholarship Council (V.Z.; B.L.), the Marie Skłodowska-Curie grant MicroSPARK (agreement no. 841466; G.K.), the Herchel Smith Funds (G.K.), the Wolfson College Junior Research Fellowship (G.K.), the BBSRC (T.P.J.K.), the Newman Foundation (T.P.J.K.), the Wellcome Trust (T.P.J.K.), and the European Research Council under the European Union’s Seventh Framework Programme (FP7/2007-2013) through the ERC grant PhysProt (agreement no. 337969; T.P.J.K.).

## Data Availability

The data that support the findings of this study are available from the corresponding author upon reasonable request.

## Supporting information

**Figure S1.**
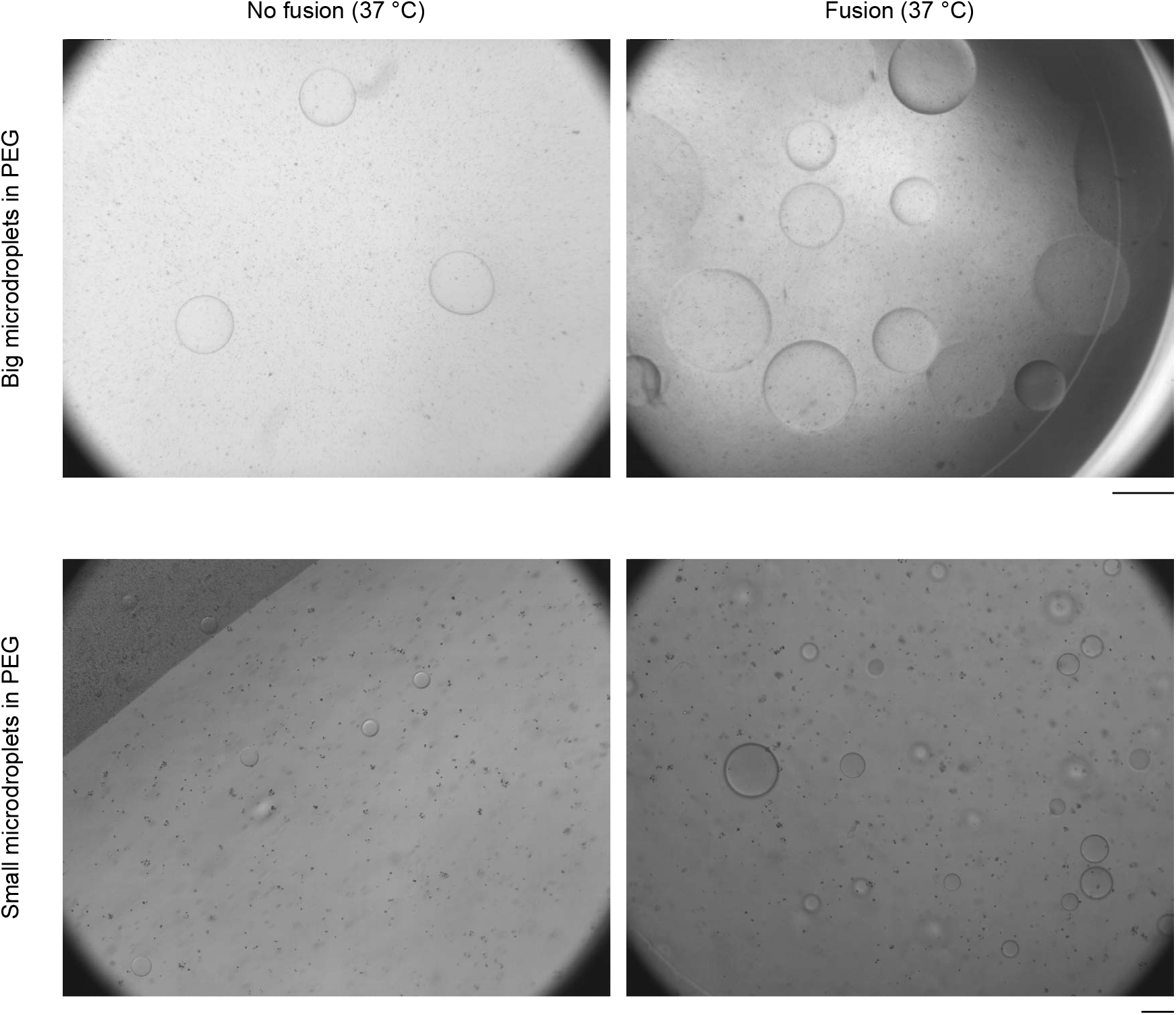
Monodisperse (left images) and polydisperse (right images) of the gelatin droplets in PEG solution at 37 °C. Big microgels (upper images; scale bar, 500 μm) and small microgels (lower images; scale bar, 100 μm) were used.

**Figure S2.**
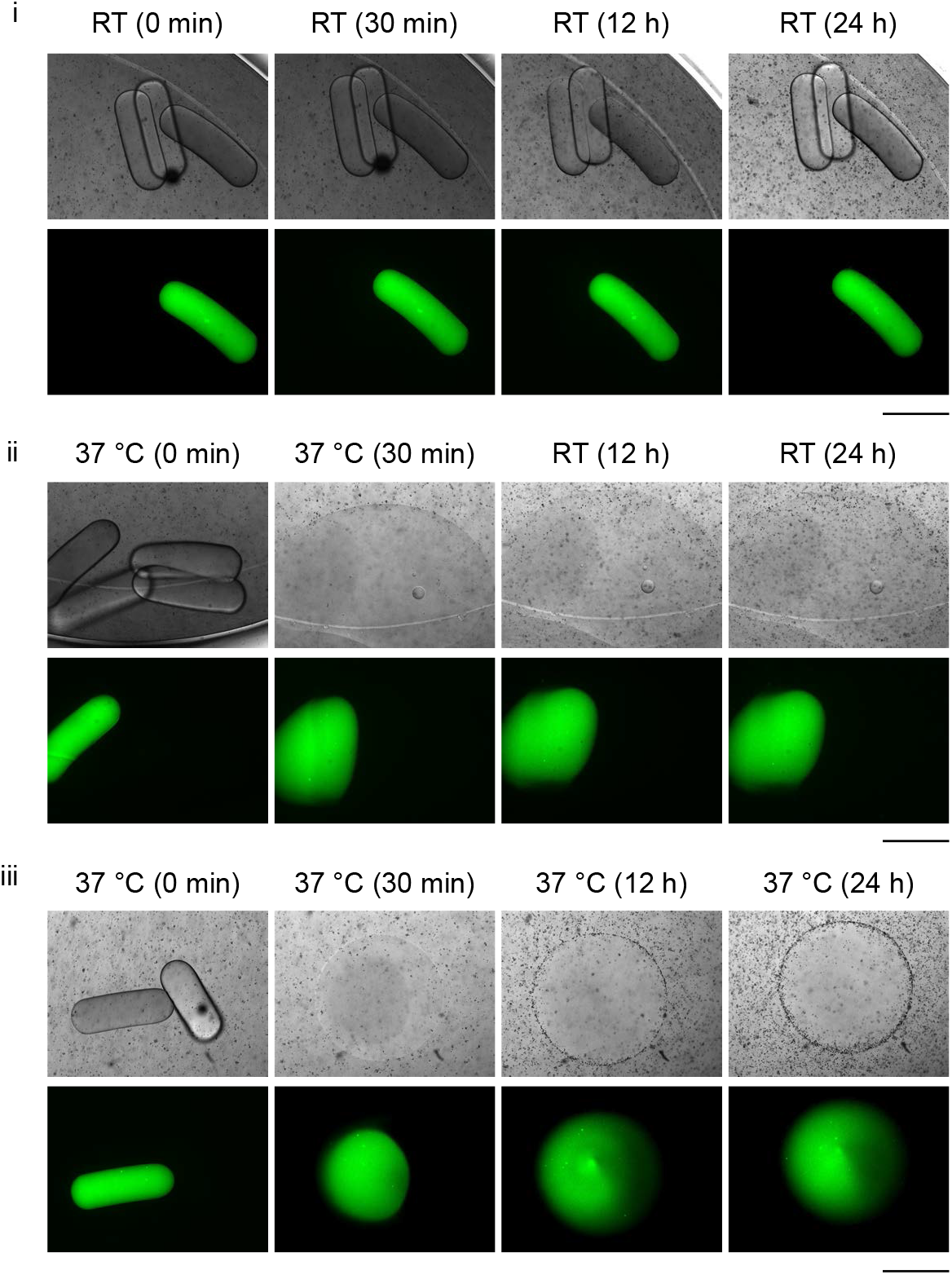
Diffusion of green nanospheres from nanosphere-laden droplets to nanosphere-free droplets. i, The microgel-PEG system was kept at RT for 24 hours. Scale bar, 500 μm. ii, The microgel-PEG system was kept at 37 °C during the first 30 minutes. Then this system was kept at RT until 24 h. Scale bar, 500 μm. iii, The microgel-PEG system was kept at 37 °C for 24 hours. Scale bar, 500 μm.

**Figure S3.**
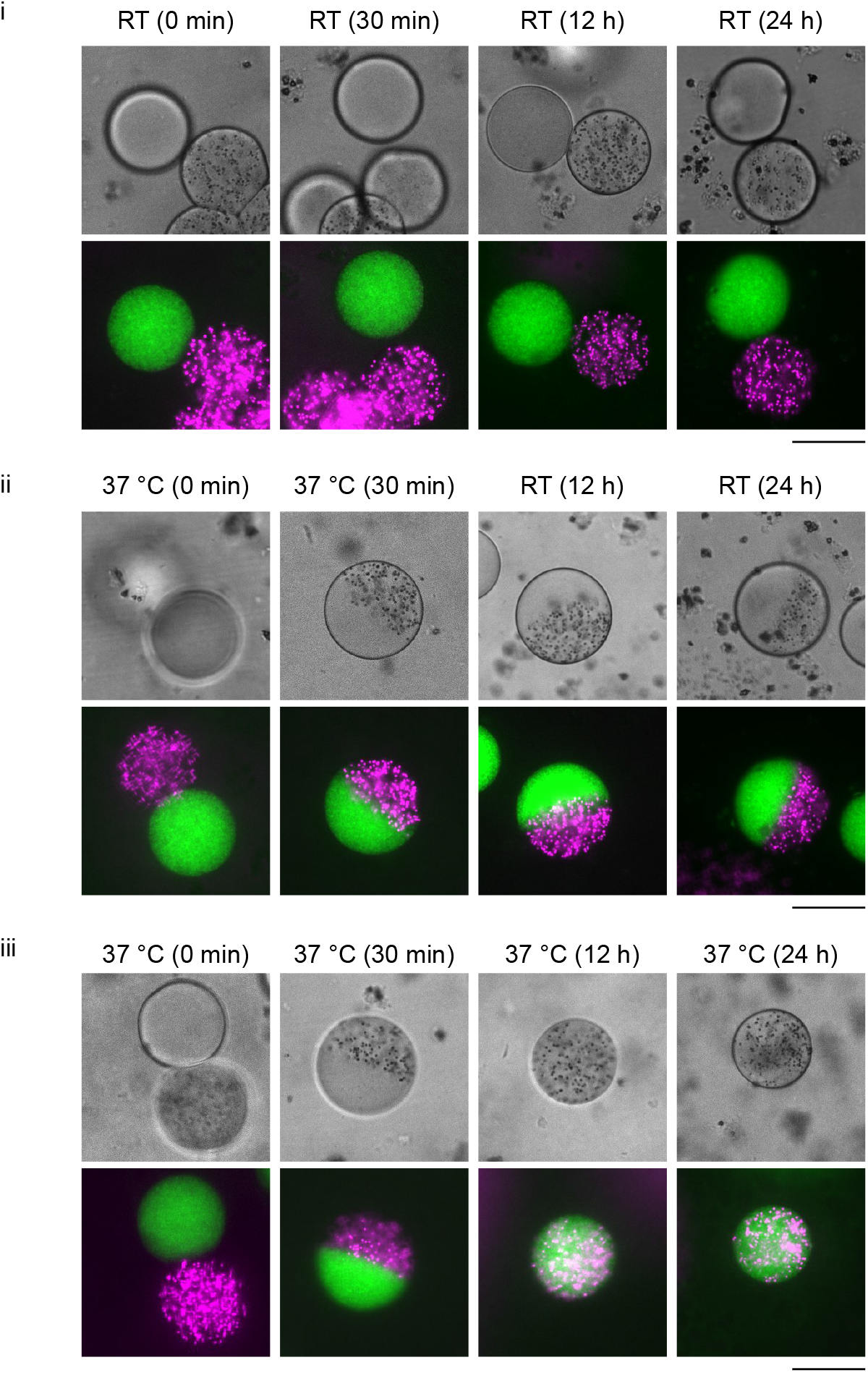
Mutual diffusion of green nanospheres and red microspheres from nano/microsphere-laden droplets. i, The microgel-PEG system was kept at RT for 24 hours. Scale bar, 50 μm. ii, The microgel-PEG system was kept at 37 °C during the first 30 minutes. Then this system was kept at RT until 24 h. Scale bar, 50 μm. iii, The microgel-PEG system was kept at 37 °C for 24 hours. Scale bar, 50 μm.

